# Dopamine Release Plateau and Outcome Signals in Dorsal Striatum Contrast with Classic Reinforcement Learning Formulations

**DOI:** 10.1101/2023.08.15.553421

**Authors:** Min Jung Kim, Daniel J. Gibson, Dan Hu, Ara Mahar, Cynthia J. Schofield, Patlapa Sompolpong, Tomoko Yoshida, Kathy T. Tran, Ann M. Graybiel

## Abstract

We recorded dopamine release signals in medial and lateral sectors of the striatum as mice learned consecutive visual cue-outcome conditioning tasks including cue association, cue discrimination, reversal, and probabilistic discrimination task versions. Dopamine release responses in medial and lateral sites exhibited learning-related changes within and across phases of acquisition. These were different for the medial and lateral sites. In neither sector could these be accounted for by classic reinforcement learning as applied to dopamine-containing neuron activity. Cue responses ranged from initial sharp peaks to modulated plateau responses. In the medial sector, outcome (reward) responses during cue conditioning were minimal or, initially, negative. By contrast, in lateral sites, strong, transient dopamine release responses occurred at both cue and outcome. Prolonged, plateau release responses to cues emerged in both regions when discriminative behavioral responses became required. In most sites, we found no evidence for a transition from outcome to cue signaling, a hallmark of temporal difference reinforcement learning as applied to midbrain dopamine activity. These findings delineate reshaping of dopamine release activity during learning and suggest that current views of reward prediction error encoding need review to accommodate distinct learning-related spatial and temporal patterns of striatal dopamine release in the dorsal striatum.

---

Pioneering work has clarified much about dopamine signaling in the brain and the remarkable relationship between this signaling and predictions of reinforcement learning algorithms. A canonical view^1^ suggests that phasic dopamine signaling acquired during learning represents a reward prediction error (RPE). This view could be formulated in terms of a temporally sequential learning-related process, by which phasic responses originally are elicited by the reward, but these responses then decline as the phasic increase in activity is transferred to the most proximal cue predictive of reward^2^. This theory-based temporal difference (TD) formulation was recognized as having clear parallels to the patterns in electrical activity exhibited by of dopamine-containing neuronal cell bodies in the midbrain recorded during learning tasks^3–7^.

Subsequent studies building on this pioneering work have demonstrated heterogeneity in dopamine responses in the substantia nigra pars compacta (SNpc) and the striatum, the two poles of the nigrostriatal tract^8–13^. The emergence of chemosensor probes to record dopamine release in the striatum, nucleus accumbens and elsewhere was transformational in opening up detailed analysis of dopamine release^14,15^. Evidence now supports the view that dopamine release in the striatum can be controlled locally and has suggested novel mechanisms of control. For example, spike activity of nigrostriatal fibers can be triggered within the striatum by cholinergic inputs acting at nicotinic acetylcholine receptors on the dopamine-containing fibers^16–18^. Dopamine release is reported to occur in waves moving across the width of the caudoputamen in ∼200 msec^19^.The release can exhibit low frequency oscillations even without task engagement, gated by extrinsic striatal afferents^20^. Topographic differences also exist. Striatal dopamine release responses can be different in different striatal sectors, prominently so between the medial and lateral regions of the caudoputamen in mice^11,21–23^. Such differences have been reported for other topographic dimensions as well^12,24–26^. The dopamine release signals can be principally related to negative as well as positive reinforcement^5,27–29^ or to non-reward parameters of movement^21,30,31^, can occur as prolonged ramping signals^32^, and can be compartmentally selective for striosome and matrix compartments of the striatum^18,33–36^. Especially for the nucleus accumbens, but also for the dorsal striatum, the relation of the release patterns to RPE-TD learning algorithms has been strongly questioned^37–39^ and strongly defended^10,23,30,37,39–47^.

We took up this issue for the dorsal striatum (caudoputamen) by training mice consecutively on a series of cue-association tasks and recording dopamine release population-level responses with dopamine sensors throughout the time that the mice were learning the tasks. Mindful of the complexities of dopamine signaling, we nevertheless looked for patterns of activity similar to those recorded in classic work on the activity of dopamine-containing cell bodies in the SNpc^3^. We found shifts in the patterning of dopamine release signals as successive versions of the cue-association tasks were acquired, and sharp differences in the dopamine release patterns between more medial and more lateral sites among the 67 mice sampled. Notably, reward outcome did not evoke transient dopamine increases in the medial sites. Over time, the cue responses declined, rather than increasing. The lateral sites did exhibit both cue and outcome responses, but they failed to exhibit a shift from primarily signaling outcome to primarily signaling cue, a canonical feature of RPE algorithms applied to the nigral dopamine system. Finally, prolonged plateau release responses to cues predicting reward emerged when the mice shifted from simple cue-association conditioning to more cognitively demanding cue discrimination conditioning, and these plateau responses appeared both medially and laterally and also were evident in somewhat modified form through the cue reversal and probabilistic reward training sessions. These discrepancies with the expectations based on cell body recordings in the dopamine-containing midbrain encourage further review of these classic algorithms.

## Results

We recorded real-time dopamine release by photometry with D1 or D2 dopamine receptor-based sensors^14,15^ placed in the medial or lateral sites of 67 mice (41 male and 26 female) that learned and performed a series of consecutively presented tasks with visually cues to instruct reward availability (**Fig. 1a-c**). These included, first, random reward presentation, and then, in succession, single-cue association conditioning, cue discrimination for two cues, reversal learning, probabilistic reward leaning, and extinction. In each task, only a single cue was presented at a time, either to the left or to the right of the mouse. For simple cue-association learning, the right or left cue, randomized across mice, was associated with reward. For the cue discrimination tasks, again only one was shown in any given trial, but two cues could be presented, one at a time. The same cue (left or right) that had predicted reward during the cue-association task was still the cue predicting reward, but its presentation alternated semi-randomly with the presentation of another cue on the other side, and it was not associated with reward. These contingencies were reversed during reversal discrimination. In the probabilistic reward task, one cue was associated with reward on 100, 75 or 50% of trials.

**Fig. 1.**
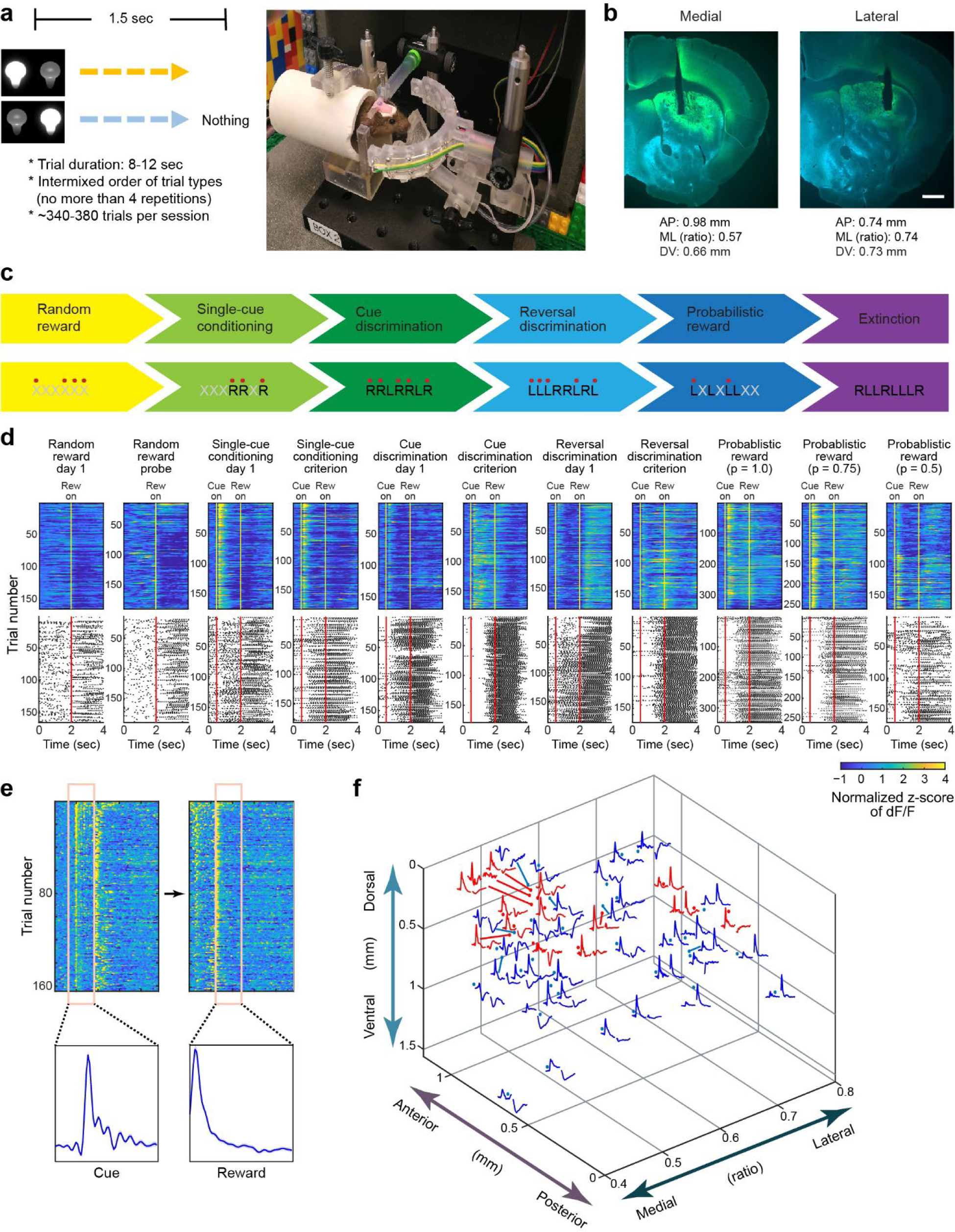
Experiment design and data analysis. **a** Head-fixed apparatus with right and left visual cues and a drop of sucrose solution as a reward (right) and trial structure of an experiment session (left). **b** Examples of histological verification of injection sites of dopamine sensors and optic probe locations in medial (left) and lateral (right) sites. **c** Experimental design showing different phases of training (each with different task contingencies, top) and examples of task events in each phase for a mouse that initially received reward following the right (R) cue (bottom). Black letters indicate cue presentation on the specified side, red dots show reward delivery, and gray Xs indicate trials in which neither cue was presented. **d** Trial-by-trial data from a single mouse, illustrating dopamine traces (top) and corresponding lick activity (bottom) for reward-predicting cues and rewards across learning sessions. See also **Fig. 2b**. Data from random reward sessions are aligned at reward delivery time (Rew on), and data from all other sessions are aligned to the cue onset (Cue on) with the 0.5-sec precue, 1.5-sec cue and 2-sec reward periods. The "Random reward probe" session is one of those inserted during later phases of training (see Methods). **e** Construction of cue and reward response traces. For each session, cue data are aligned to the onset of cue presentation, and reward data are aligned to the first lick after reward delivery. Note that the examples show two different sessions. **f** 3D reconstruction of dopamine responses to cues and rewards according to histological confirmations of probe locations in standardized coordinates (see Methods). Dopamine release was recorded with D1R-based (red) and D2R-based (blue) sensors during the first session of the single-cue conditioning.

Each mouse was implanted with a single optic probe in either the medial sector (36 mice) or lateral sector (31 mice) of the caudoputamen of the right hemisphere (**Fig. 1b, f**). This unilateral, single-probe recording protocol was chosen to minimize potential damage to the striatum and damage to the overlying neocortex due to the insertion of the probe, important given the extended periods of chronic recording required for the mice to complete the six different tasks. **Fig. 1d** illustrates for one mouse the dopamine release patterns and corresponding licking activity throughout learning, scored according to the method indicated in **Fig. 1e** and Methods.

Initial random reward sessions were given to allow assessment of dopamine release responses to unpredicted rewards. Reward-evoked dopamine in the medial and lateral sites (**Fig. 2a**) exhibited marked differences. In the medial sites, the responses to randomly delivered rewards at first dipped below baseline release levels, then flattened to low positive levels. By contrast, the release signals evoked in the lateral sites were at first small negative-positive blips that soon developed into strong positive, transient responses. For the full sequence of task versions that followed, random reward sessions were inserted every 9 or 10 sessions to determine whether the release responses were affected by the progressive learning stages (Methods). We did not observe different results across the data sets acquired with the D1- or D2-based probes (data not shown), and we merged these for the analyses.

**Fig. 2.**
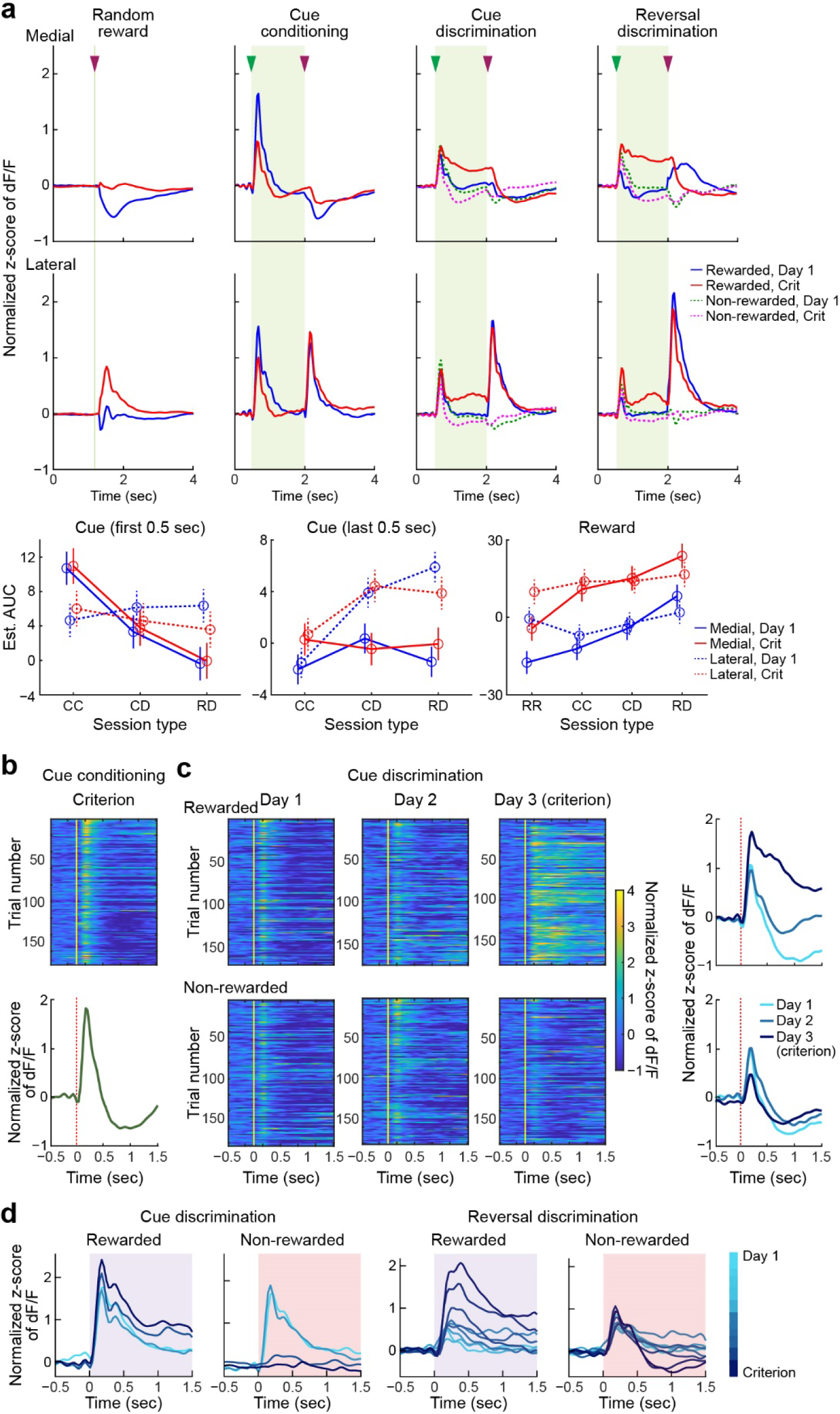
Distinctive characteristics of dopamine release signals in dorsal striatal subregions evoked during a series of conditioning tasks. **a** Learning-related effects on the dopamine responses to the cue (green arrowheads) and reward (purple arrowheads) in medial (top) and lateral (middle) sites. Averaged traces for all mice of the first (Day 1) and criterion (Crit) sessions for each session type are shown. Data from random reward sessions, which included those inserted late in training, are aligned with reward delivery at t = 1.0 sec. Plots for other session types span 0.5-sec precue, 1.5-sec cue (light green shade) and 2-sec reward periods. For discrimination learning, average traces of non-rewarded trials are shown with dotted lines. Summary graphs of post-estimation by GLMM shows estimated fit and 95% confidence intervals for dopamine response in early (first 0.5 sec) and late (last 0.5 sec) cue periods and reward period of random reward (RR), cue conditioning (CC), cue discrimination (CD) and reversal discrimination (RD) sessions. Dopamine responses were quantified as AUC of the dF/F data (see Methods) for the rewarded trials. **b, c** Transition from initial cue-association (**b**) to cue discrimination (**c**) training shown for a single mouse. In **c**, dopamine response in rewarded (top) and non-rewarded trials are shown. Vertical lines indicate cue onset. Dopamine traces aligned as in **Fig. 1d and e** illustrate responses in consecutive sessions from last cue conditioning (**b**, bottom) to first three cue discrimination sessions (**c**, right), illustrating gradual emergence of prolonged plateau dopamine release during cue presentation period in rewarded trials of cue discrimination sessions. Color scale shows z-scores of dF/F, which ranged from −1 to +4. Color-coded line plots on right. **d** Superimposed session-averaged dopamine release in response to rewarded and non-rewarded cue onsets recorded in a mouse during all training sessions, from day 1 (light blue) to criterion (dark blue), of cue (2 left panels) and reversal (2 right panels) discrimination training. Shaded purple and pink boxes indicate 1.5-sec cue period.

In the first cue conditioning sessions, in which the mice learned to associate the cue with reward, the medial sites and lateral sites again exhibited clearly different release dynamics (**Figs. 1f** and **2**). In the medial sites, strong positive transient dopamine release responses occurred at cue, but responses at outcome (registered as the first lick after the reward delivery) were at first negative (below baseline) and then nil or weakly positive. The lateral sites, by contrast, exhibited strong phasic responses at outcome as well as at cue onset, and these responses remained stable throughout the sessions. Outcome signaling thus was exhibited in the lateral sites but was weak or negative-going in the medial sites. In both the medial and lateral regions, the cue responses were diminished in amplitude by the late training sessions, not enhanced as expected from RL-RPE accounts. Most notably, we were unable to identify a ‘transfer’ of the dopamine signaling from outcome to reward-predictive cues as suggested by classical work based on recordings in the dopamine-containing cells of the SNpc^1,3^.

We searched for such dynamics by continuing the training to require the mice to discriminate which of two cues was associated with reward. The originally rewarded cue was now randomly alternated with a second cue, which appeared on the opposite side and did not predict reward. Again, the dopamine responses recorded at outcome were nearly absent medially and in the lateral sites did not decline with training, in sharp contrast to the predictions of classical RPE (**Fig. 2a**). Remarkably, both medial and lateral cue responses became sustained. They persisted during most or all of the 1.5-sec long cue period as learning proceeded, then fell at outcome in the medial sites and rose transiently at this cue-off/outcome-on time in the lateral sites. These plateaus emerged during the first days of discrimination training (**Fig. 2b, c**).

The development of the plateau-like response to the reward-predicting cue was not the product of averaging across mice; it could be seen in individual mice. In the example shown in **Fig. 2d**, at the start of cue discrimination training, both cues produced a slowly decaying dopamine response that extended well past the initial peak. As training proceeded, the response to the reward-predicting cue became larger and more sustained, whereas the response to the non-reward-predicting cue was nearly abolished by the third session. A similar process occurred during cue reversal training (see following), and by the time the mouse reached criterion, the long latency component of the dopamine response to the non-rewarded cue showed an anti-plateau, i.e., a sustained drop below baseline.

After each mouse reached criterion for cue discrimination, the mouse was trained on a reversal discrimination task that required the mouse to learn that the formerly rewarded cue was now the non-rewarded cue. This task version again required cue discrimination learning for success in obtaining reward. Prolonged plateau dopamine release responses to cue occurred in both medial sites and lateral sites (**Fig. 2a**). In the lateral sites, outcome signaling was strong and steady, and there was a small increase in the cue response. These results again were not anticipated on the basis of RPE algorithms. More medially, a positive outcome signal emerged for the first time, but then waned with exposure across trials. This brief positive signal, unique to the beginning of reversal training, suggests that in some medial sites, outcome responses might be associated with reversing the previously learned association, a pattern compatible with dopamine serving as an RPE signal.

To characterize further the relationships between dopamine, learning, and RPE, we introduced a task version with probabilistic reward sessions, in which one cue (the same one that was rewarded in the preceding reversal training) always signaled potential reward, but with varying probabilities (**Fig. 3a, b**). The responses in the medial sites, instead of being dominated by a single sustained plateau, now had two components: an early strong transient followed by a much lower amplitude sustained plateau response that was greater than baseline for the high-probability conditions, but essentially zero for the 50% rewarded condition. There were dips at the no-reward outcomes, but little or no response to reward outcomes. By contrast, the lateral sites again exhibited stability in their strong dopamine release transients both at cue and at positive outcomes, with dips for non-rewarded trials. Notably, the magnitude of the lateral outcome responses clearly scaled with probability of reward, systematically increasing with lower probabilities of reward during the probabilistic reward sessions, as though scaling with uncertainty (largest response at 50% probability). These response patterns are consistent with an RPE interpretation, but once more, there was no sign of response transfer from outcome to cue.

**Fig. 3.**
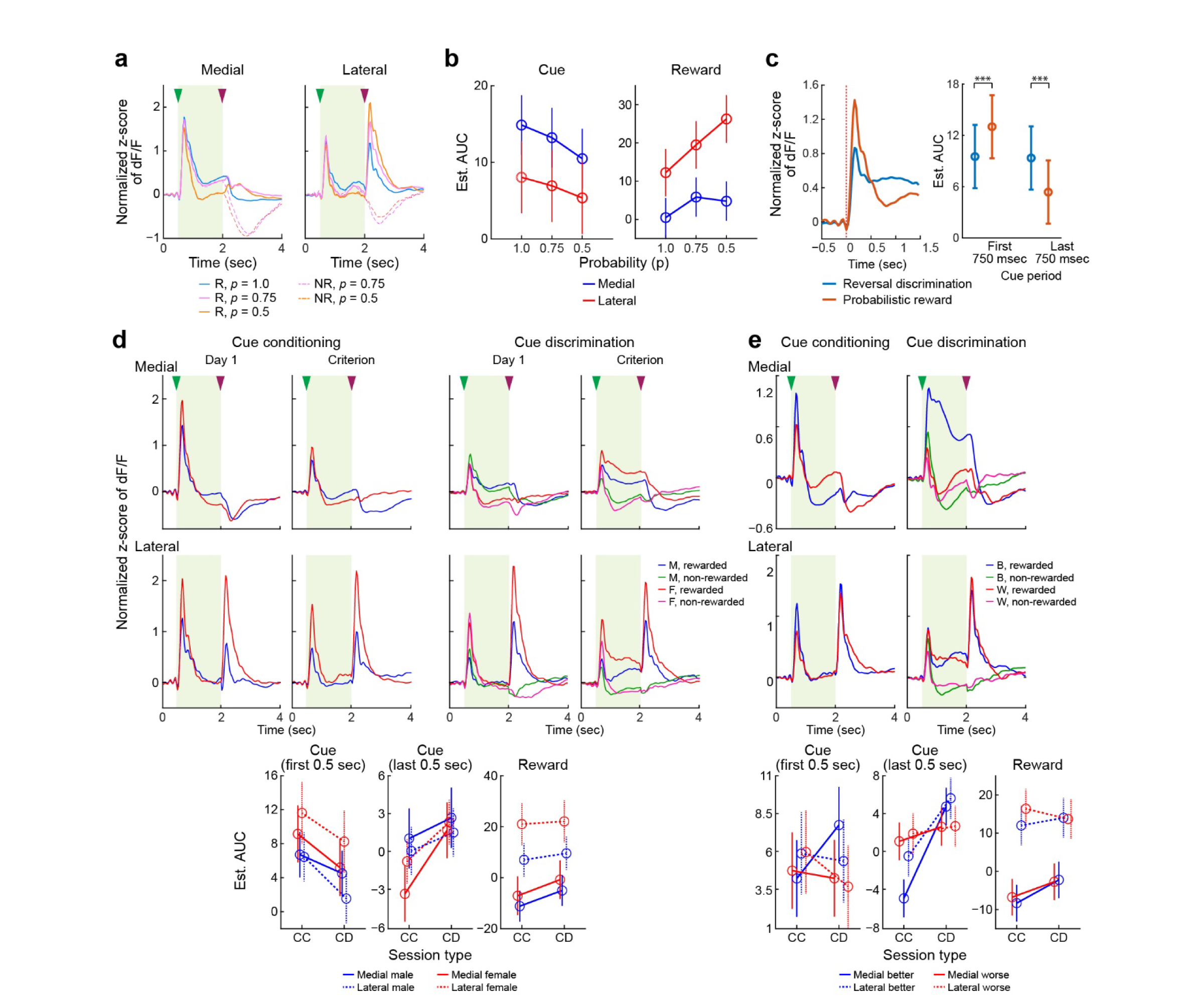
Effects on dopamine response of reward probability, biological sex, and discrimination performance. **a** Dopamine response to cue and reward during the probabilistic reward sessions recorded in the medial and lateral regions, shown separately for rewarded (R) and non-rewarded (NR) trials with different reward probabilities (*p* = 0.5, 0.75 and 1.0).. Green and purple arrowheads represent, respectively, cue and reward onsets. **b** Dopamine response AUC values for cue and reward periods in sessions with different reward probabilities. The marginal means and 95% confidence intervals for the cue and reward response are shown, according to the different reward probabilities, in relation to the recorded sites. **c.** Left: Average dopamine response to reward cue during the last reversal discrimination session and the first probabilistic reward (PR) session. Right: The post-estimation of GLMM of dopamine signal for the reward cue showing significant differences in early (the first half) and late (the second half) cue periods between reward discrimination and probabilistic reward sessions. \*\*\**p* < 0.0001. **d** Average dopamine traces recorded during the first (Day 1) and last (Criterion) sessions of cue conditioning (CC) and cue discrimination (CD) learning at medial (top) and lateral (middle) sites. Traces are color-coded for male (M) and female (F) mice, and rewarded and non-rewarded trials. Bottom row depicts post-hoc estimation of dopamine response AUC for first and last third of cue period, and entire reward period, calculated from the rewarded trial data. Day 1 and Criterion sessions were combined for CC, and separately combined for CD, for each AUC period. Error bars as in **b**. **e** Average dopamine responses recorded in the last (Criterion) sessions for cue conditioning and discrimination learning plotted for better than median (B) and worse than median (W) performers, and for rewarded and non-rewarded trials. Post-estimation of response. Bottom row: AUC for cue and reward periods from the rewarded trial data, as in **d**. Error bars as in **a**.

At the transition from cue reversal learning to 100% probabilistic reward sessions, the average dopamine response changed in shape (**Fig. 3c**). The initial peak increased in height by nearly twofold, and the plateau height dropped by a similar amount. Based on the GLMM interaction model and the pairwise comparisons, responses to the rewarded cue changed significantly from reversal discrimination to probabilistic reward sessions in both the initial and later cue periods (*p* < 0.0001).

To look for possible factors contributing to the cue and outcome responses, we examined signal differences based on the animal’s sex (**Fig. 3d**) and on the basis of their performance levels in the task versions (**Fig. 3e**). Across task versions, dopamine release level changes (both positive and negative) were larger for the females than for the males, but were similar in pattern, thus exaggerating the contrast between the medial and lateral dopamine release patterns in females (**Fig. 3d**). We also plotted the trial-averaged dopamine responses for the first and last sessions of cue conditioning and cue discrimination, averaged over the group of mice that performed better than median (better performers), and separately averaged over those that performed worse (worse performers) (**Fig. 3e**). In medial sites, dopamine responses were strikingly different for the better and the poorer performers in both the cue conditioning and cue discrimination tasks, and they exhibited a strong difference between rewarded and non-rewarded trials in cue discrimination that was absent from the worse performers’ patterns. By contrast, the dopamine responses at lateral sites were similar regardless of performance level across all task conditions.

The amplitudes of the plateau response at cue onset generally followed the levels of task acquisition for many mice (**Fig. 4a**), high for most of the good learners, less prominent for the middling learners, and not detectable in the mice that did not learn well or at all. The same trends were present for reversal discrimination responses to the previously non-rewarded cue (**Fig. 4b**). Each of the data sets was accompanied by analysis of the responses that the mice made to the non-rewarded cue. Many of the proficient and moderately good learners developed brief transient responses to the non-reward-predicting cue, but those peaks were not followed by plateaus. Correlation analysis showed that among mice that reached the learning criterion in the cue discrimination task, there was a highly significant (p = 0.002) correlation between learning index and the difference in area under the curve (AUC) of the dopamine responses to reward-predicting and non-reward-predicting cues (**Fig. 4c**). A similar result was obtained for the cue reversal task (p = 0.02; **Fig. 4d**).

**Fig. 4.**
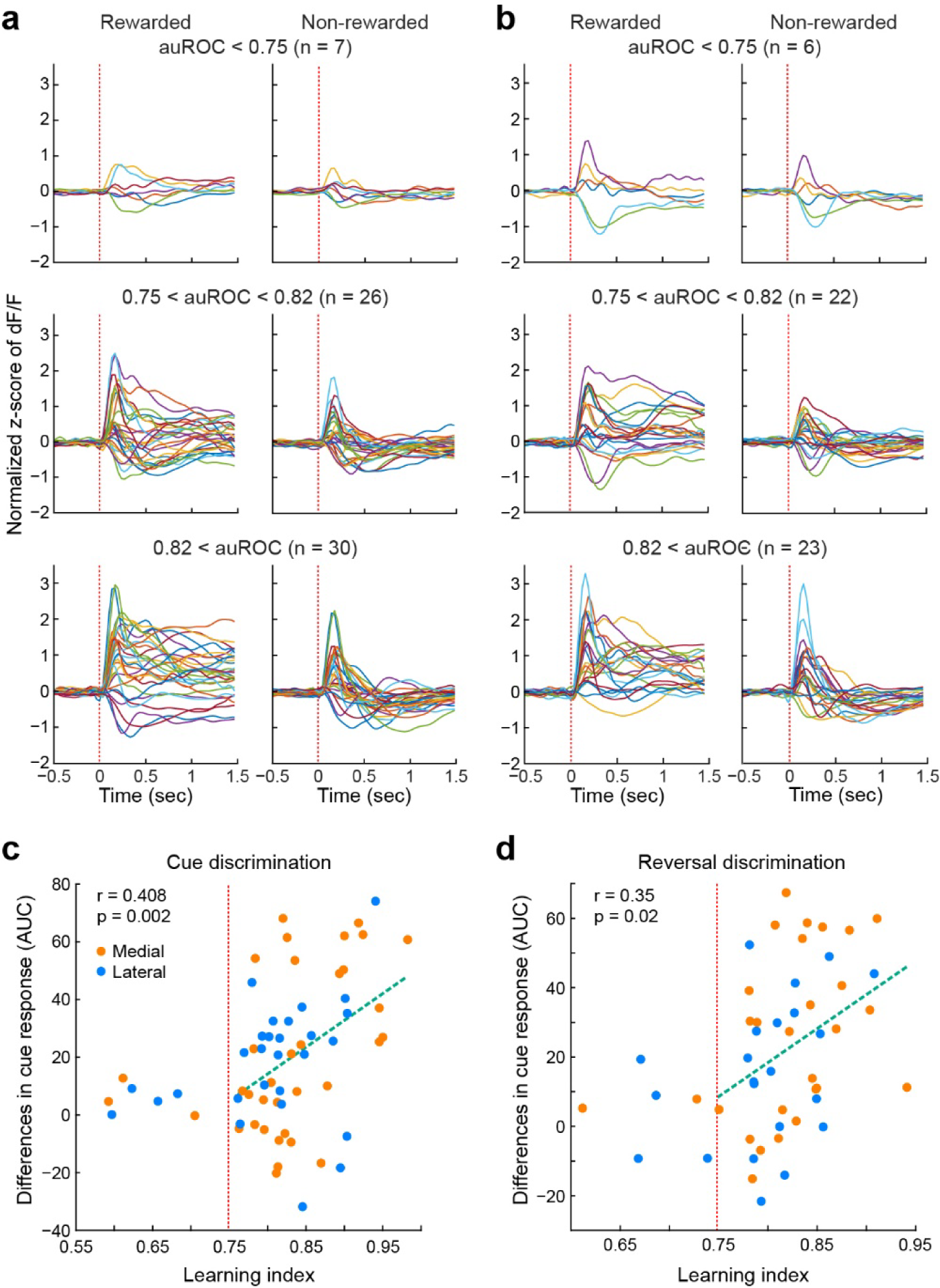
Dopamine plateau and discrimination learning. **a, b** Dopamine responses to cues predictive of reward (left) or no reward (right) shown for mice according to their final performance rates (top to bottom, poor to best learners) during cue discrimination (**a**) and reversal discrimination (**b**) sessions. Each trace represents one mouse. Vertical line shows time of cue onset. **c, d** Scatter plots of learning index and differential cue responses between rewarded and non-rewarded trials during the last session of cue discrimination (**c**) and reversal discrimination (**d**). Each point represents one mouse (orange for medial sites, blue for lateral). Vertical red line shows learning criterion.

The existence of a variable plateau-like component in the dopamine responses was further confirmed by performing principal component analysis (PCA) on the cue discrimination data (**Fig. 5**) to identify correlated components of variance across waveforms, sorted in order of decreasing variance explained. Each mouse was represented in the PCA by the response waveform during the cue period averaged over all rewarded trials in that mouse’s first ("Day 1") and final ("Criterion") sessions of discrimination training, and an analogous averaged waveform for the reward period. **Fig. 5a** shows the waveforms for the cue period, with the grand average across all mice shown in black, and the first three principal components of the variance (PCs) in different colors. The average waveform has a prominent plateau at about half the amplitude of the initial peak. The shape of the first PC, in red, indicates that the height of the plateau could vary independently from the height of the initial peak, and the higher the plateau was, the more it tended to decay slowly over time. The first PC for this data set accounted for the vast majority of the variance across mice, and therefore across recording locations (**Fig. 5a**, right). For the reward period (**Fig. 5b**), PC1 again accounted for more than half the variance in the waveforms and demonstrated that a slowly decaying plateau is the chief type of variation across different waveforms. There was a slight correlation between plateau height and peak height, but the prominence of the peak in both PC2 and PC3 indicated that there was also considerable independence between peak height and plateau height.

**Fig. 5.**
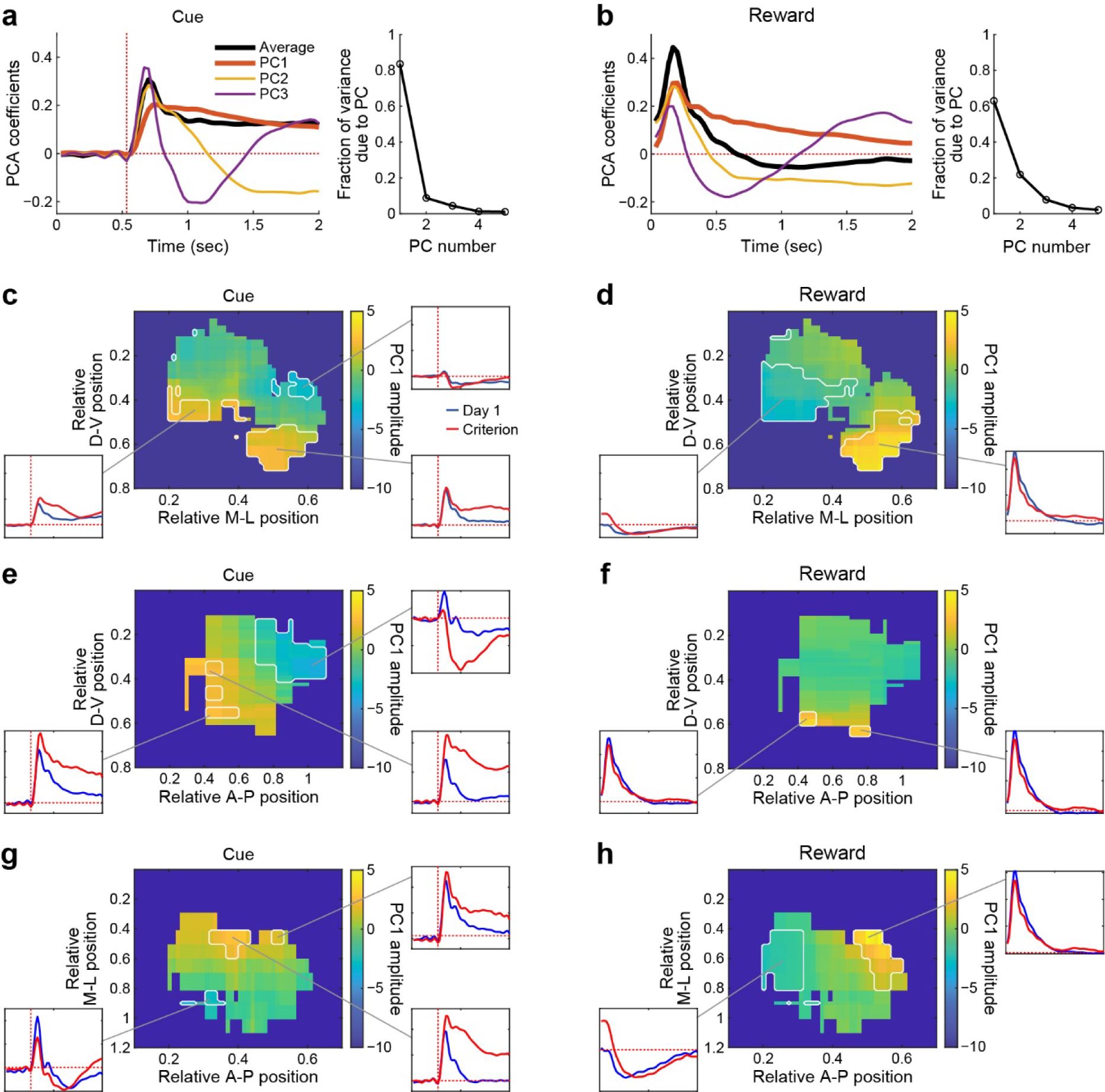
Amplitude of first principal component of fluorescence waveform as a function of probe position in striatum after discrimination learning. **a** Left: Average response waveform across all mice in the last session of discrimination training (i.e., the "learned" state), and first three principal component waveforms, during the cue period. Average waveform has been normalized to unit magnitude to match the scale of the principal component waveforms. Vertical dotted line indicates cue onset. Right: Fraction of waveform variance across mice explained by each of the first five principal components. **b** Same as **a**, but for reward period, starting at reward onset. **c** Anatomical distribution in coronal plane of first principal component amplitudes during cue period. Color scale shows average amplitude of PC1 across all recording sites that were within 8 spatial bins of each point. Only points that had at least 3 recording sites contributing to the average are plotted; other points are shown in dark blue. White outlines show regions that were significantly different from median (*p* < 0.05, two-tailed bootstrap). Inset plots show fluorescence waveforms recorded in the first (Day 1) and last (Criterion) sessions of discrimination training, and averaged across probes that contributed to the average in the middle of the indicated region. All insets are shown with z-score normalized dF/F ranging from −0.5 to 2.7 on the vertical axis, and time spanning 2 sec on the horizontal axis. Dotted vertical line indicates cue onset, dotted horizontal line marks dF/F = 0. **d** Same as c, but for responses in reward period. Insets do not include the time of cue onset, but start at reward onset as in **b**. **e, f** Same as **c** and **d**, but projected onto sagittal plane instead of coronal. **g, f** Same as **c** and **d**, but projected onto horizontal plane instead of coronal.

It was clear by eye that the prominence of the plateau components following both cue onset and reward delivery differed across different subregions of the striatum. We therefore constructed spatially smoothed maps showing the variation in average amplitude from multiple probes that were in the same vicinity (see Methods). Such maps are shown for PC1 amplitude projected onto the three cardinal anatomical planes, aligned at cue onset (**Fig. 5c, e, g**) and reward delivery (**Fig. 5d, f, h**), with shades of yellow representing high values of PC1 amplitude, and shades of blue or green representing low or negative amplitudes. The changes across learning stages from early training to acquisition are equally striking (**Supplementary Fig. 1**). The maps in all three anatomical planes exhibited substantial spatial variation, so that across the 3D expanse of the striatum, there were districts with strong changes in plateau levels and learning-related development and others where they were not so prominent. The PC1 component of the response waveforms was significantly higher in the ventrolateral region for both cue and reward periods, but in the ventromedial region, it was significantly lower in the reward period and significantly higher in the cue period. The distribution of dopamine response waveforms were quite different following cue onset and reward delivery.

Dips in dopamine release going below baseline levels occurred early on in training sessions at the end of the cue period (beginning of the availability of reward). These were also present in the random reward sessions. We therefore asked whether the licking patterns themselves could have been important in shaping the release response profiles as the mice adjusted these patterns and formed stereotyped licking patterns toward cues and reward. We aligned the dopamine signals relative not only to the first lick after reward availability (end of cue), as in **Fig. 2**, but also to the spontaneous, un-cued licks that occurred during the inter-trial intervals, identified as having at least a 0.5-sec period without licks prior to the spontaneous lick (**Fig. 6a**); and also to the first lick after cue-onset (anticipatory licking; **Fig. 6b**). In all instances, the dopamine response was negative, occurred at both lateral and medial sites, and was generally greater medially. The magnitude of the negativity waned as sessions continued (**Fig. 6b**). In the early cue conditioning sessions, there was very high dopamine release early in the trials, followed by very large reductions at the first lick after cue onset, both medially and laterally. The high dopamine release diminished during single-cue association training, resulting in a smaller decrease at first lick. In sharp contrast, in both the cue discrimination and cue reversal discrimination tasks, there was little change in dopamine release (**Fig. 6b**; note differences in vertical scales).

**Fig. 6.**
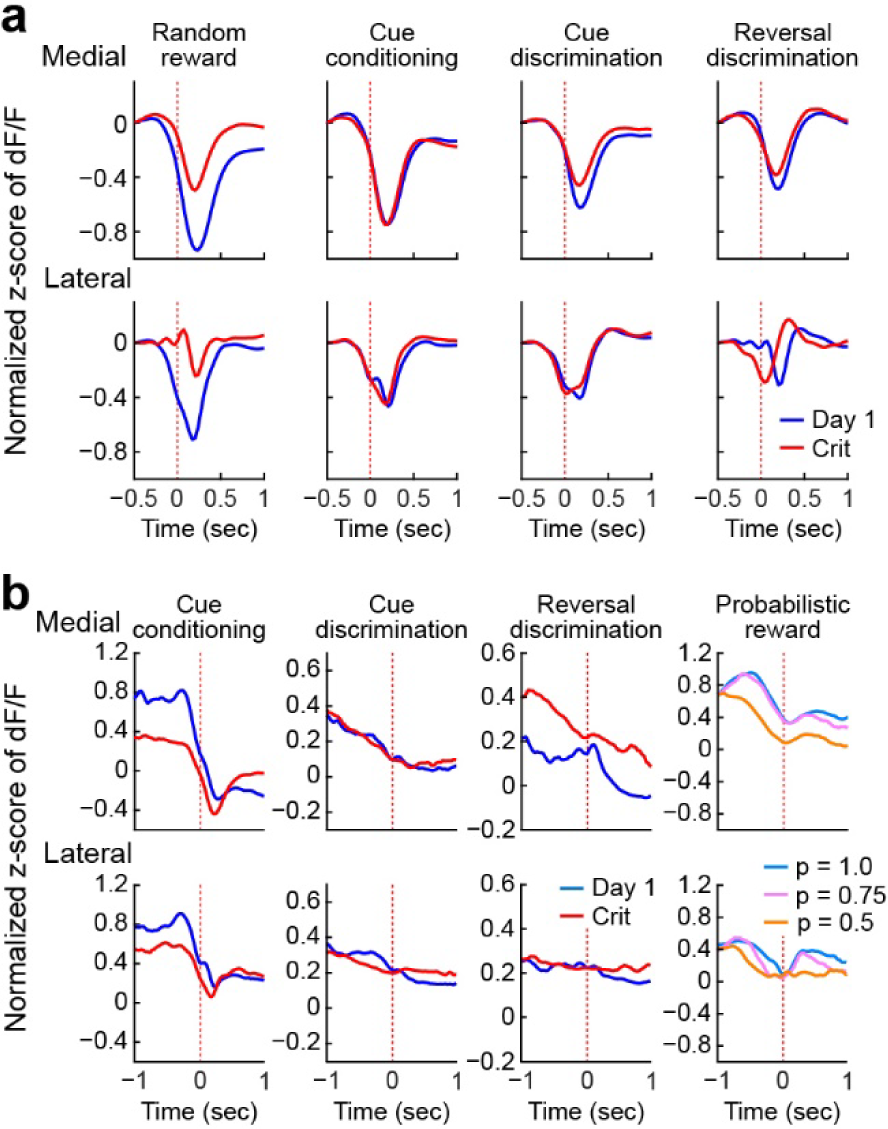
Average dopamine release in relation to licks. **a** Averaged dF/F dopamine traces aligned to spontaneous, out-of-task licks (red vertical lines) recorded in the first (Day 1) and last (Crit) sessions of each phase of training. **b** Averaged dF/F traces aligned to first lick after cue onset. Color code for last column represents the probability of receiving reward after presentation of the reward cue.

## Discussion

We demonstrate here that dopamine release profiles in the dorsal striatum undergo dynamic learning-related changes with transient increases and dips, as are well known, but also prolonged plateau release responses. These different release profiles were selective for striatal region and were selective for different versions of associative cue-outcome conditioning. We chose to use these simple conditioning tasks to connect with classic evidence for RPE reinforcement learning profiles of neurons in nigral dopamine-containing neurons^1^. During single cue association sessions, large transient increases in dopamine release occurred in the medial sites in response to the cue predictive of reward, but the medial sites lacked increased release responses at outcome (reward). Lateral sites, by contrast, exhibited strong phasic increases in release both at cue and at positive outcome, and contrary to expectations based on RPE, their response to the cue decreased with learning and their response to reward delivery remained about the same across training. Remarkably, when the mice proceeded from single cue association to cue discrimination training, prolonged plateau responses to the cues appeared in both medial sites and lateral sites. These plateaus extended throughout the cue period during cue discrimination, reversal discrimination, and probabilistic reward training, with slightly different forms suggesting that a two-component initial phasic increase carried into a plateau release of dopamine largely continuing through the cue presentation period.

We followed these across training phases and found that they were most pronounced in mice with the most proficient performance and were nearly absent in the slow or non-learners. The sources influencing these learning-related plateaus were not identified; but it is likely that in some way, the presence of an alternative cue, even if not immediately visible at the time of a response and not indicative of reward availability, could induce a network state change leading to the tendency for an extended cue response. It was only for outcome signals recorded in lateral sites that we could detect systematic changes related to changing probabilities of reward. Despite these uncertainties, our observations introduce evidence for dopamine plateau responses as learning-related features to add to transient and ramping responses formerly reported, and raise new questions about RPE encoding by the striatum during learning^39^.

In a notable deviation from classic RPE models, we did not observe as learning proceeded a transfer of the dopamine responses from the time of outcome to the time of the outcome-predictive cues. Instead, in all versions of the task, cues were signaled by dopamine release in both medial sites and lateral sites throughout training, and outcome was accompanied in the lateral sites by dopamine release throughout training. Temporal transfer and the development of RPE signals have been questioned before (reviewed in Refs 37,39 and 40), but in recent work by Watabe-Uchida and colleagues^40,41^, both RPE phenomena have been shown for the ventral striatum/nucleus accumbens and its ventral tegmental area afferent dopamine-containing neurons both at a population level and at the level of single cells. Our findings for the dorsal striatum suggest that further refinement and extension of reinforcement learning algorithms is needed to account for spatiotemporal dopamine release dynamics in the dorsal striatum, as here represented by recordings in central-medial sites and central-lateral sites, and also for the different release dynamics recorded during multiple phases of cue-association conditioning.

The transfer of dopamine signaling from outcome reinforcer to the most proximal predictor of that outcome is a central feature of RPE algorithms as applied to neural activity in the dopamine system. How could the lack of such transfer in our mice be accounted for? One possibility is that we missed this outcome-to-cue transfer in our dopamine release recordings because these were population measurements with incomplete coverage of the dorsal striatum, hiding sub-populations conforming to the RPE predictions or sub-populations not in the range of our probes. This possibility is clearly high on the list of issues needing further testing, but it does nothing to account for the behavior of other aspects of the findings that do conform to RPE. An alternative possibility is that local circuits in the striatum affected by top-down signals from the thalamus and neocortex or elsewhere could modify the striatal firing of nigral dopamine-containing axons/terminals to block outcome signaling in the medial sites in our experiments. Intrastriatal modulation of dopamine release by cholinergic interneurons, proposed for many years based on striatal pharmacology^48–50^, can change their activity during learning^4^. For example, some cholinergic inputs, some likely from these interneurons, generate action potentials in intrastriatal dopamine fibers far from their cell bodies^16^. Further, oscillatory local field potentials can accompany and even modulate activity^39,51–53^. We did not monitor this activity. Yet another possibly is suggested by the report by Hamid et al.^19^ that dopamine release signals in the caudoputamen occurs in mediolateral and lateromedial waves, moving during Pavlovian conditioning from lateral to medial at rates of about 200 msec per transit. This activity was mainly recorded in the context of heavy damage to the overlying neocortex, which we tried to avoid here by using a single probe per mouse, but it could potentially comprise a scanning mechanism sensitive to such signals as we report here, imposed by yet unknown afferent or intrastriatal circuit elements. Further dynamics of intraneuronal networks in the striatum surely must contribute. Across all these possibilities, at the population level, the patterns of transient and plateau release of dopamine in the dorsal striatum, as measured here with two different sensor types, was quite distinct from, and difficult to align with, classical RPE algorithms.

Each district of the striatum, as each area of the neocortex, likely uses and encodes different aspects of reinforcement along a nuanced scale from appetitive to aversive reinforcement options. The striatal processing surely must involve much higher-dimensional algorithms than one just dealing with expected appetitive to aversive value, or only RPE; and different sectors of the striatum and corresponding corticostriatal circuits are engaged by different components of task execution^54^. Our recordings were limited to central-medial and central-lateral sites in the dorsal striatum, and thus did not fully span the striatum as now can be done with emerging methods^55^. For dopamine release recorded in the ventral striatum/nucleus accumbens in a series of conditioning tasks, Jeong et al.^37^ have found inconsistencies between dopamine release signals there and the predictions of RPE formulations. These favor what the authors term as a retrospective causal learning algorithm. The recordings made by Jeong et al., like our recordings, were made from dopamine-containing axons, not from dopamine cell bodies as in the original studies linking dopamine dynamics to RPE. These discrepancies could be accounted for according the Uchida and Watabe-Uchida and Uchida groups (e.g., Ref 40; see also Ref 39).

Here, we have shown discrepancies with RPE formulations in both space and time. In summary, (1) there are spatially restricted representations of outcome, with outcome signals minimal or even negative in most rostral and medial regions that we explored; (2) a fundamental requirement of RPE theory, i.e., that responses move from outcome to the most proximal cue predicting reward, was not reflected in the recordings that we made in either the medial sites or the lateral sites; and (3) transient dopamine releases at cues predictive of reward were present for simple cue-association paradigms, but for the training paradigms engaging greater cognitive load, these were replaced by complex responses incorporating prolonged plateau responses with increasing, decreasing, or biphasic components superimposed. Understanding the roots of these disparities could help to uncover the remarkable functional range of dopamine-based systems in modulating adaptive behavior.

## Methods

### Animals

All experimental procedures were performed on 4-6 month-old wild type mice and F1 hybrids on C57BL/6J (Jackson Laboratory, strain ID #: 000664) and FVB (Taconic, model #FVB) background with the approval of the Committee on Animal Care at the Massachusetts Institute of Technology (MIT). F1 hybrids were produced from FVB mice in which *Pde6brd1* and *Disc1* were bred out (‘corrected FVB’). Mice were group-housed separated by sex at 25°C, 50% humidity with a 12:12 hr light/dark cycle until the intracranial injection of sensors and optic fiber and head bar implantation. Subsequently, mice were single-housed in home-cage environment enriched by addition of eco-bedding, nestlets and a PVC tube matching their body length as a play tunnel and then a body case during subsequent experimental sessions. Training sessions were conducted during the light cycle, 3-6 hr after the daylight cycle switch. Mice were placed for at least 20 min in the experimental room after transport from the vivarium before testing.

### Mouse preparation

Prior to daily training, mice (n = 80 prepared in total, 67 tested, 13 mice later excluded due to 2 misplaced probes, 3 implant detachments, 2 mouse illnesses, and 6 unidentifiable probe locations) underwent stereotaxic surgery twice for virally mediated injection of a single dopamine sensor, either GRAB_DA2m (AAV9-hSyn-DA4.4), GRAB_DA3m (AAV9-hSyn-DA3m), or dLight1.3b (AAV5-CAG-dLight1.3b), followed by optic fiber and head-bar installation one week later. Mice deeply anaesthetized with isoflurane (1-2% on oxygen flow rate of 0.4 L/min) were mounted on a stereotaxic apparatus, and were injected subcutaneously with buprenorphine (2 mg/kg) and meloxicam (2 mg/kg) as pre-surgical analgesics, and for 3 days post-surgically as needed. The skin covering the skull was incised and a burr hole was made to place an injection needle to carry the viral construct to the target site in the right hemisphere (AP: +1.0 mm, ML: +1.7 mm, DV: −3.1 mm from bregma). A 0.5 µL aliquot of viral construct was administered a rate of 0.05 µL/min. The injection needle was left in place for 5-10 min after completion of the injection and then was slowly removed from the brain. The burr hole was filled with bone wax and the overlying skin was sutured shut. One week to 10 days later, mice were anesthetized as before, mounted in the stereotaxic machine, and the burr hole exposed and enlarged medially. The exposed skull was cleaned with cotton swabs dampened in 3% hydrogen peroxide, scarified using the tip of a surgical scalpel to enhance subsequent bonding of bone cement. The optic fiber was inserted to the target position (AP: +1.0 mm, ML: +1.5 mm, DV: −2.7 mm from bregma for medial sites; AP: +1.0 mm, ML: +1.9 mm, DV: −3.0 mm from bregma for lateral sites), the burr hole opening was filled with a small amount of petroleum jelly, a thin layer of Metabond was applied to the exposed skull and to the bottom face of optic fiber ferrule. A 3D-printed head bar (3.2 mm (W) x 3.2 mm (H) x 25.5 mm (L); weight 0.25 g) was positioned −2 mm posterior from the lambda and securely cemented onto the Metabond treated skull. Mice were allowed to recover for at least 3 weeks and left undisturbed except for regular animal husbandry care.

Thereafter, each mouse was mounted in the head-fixed recording chamber, and spontaneous dopamine signals were collected for an hour to check signal quality. Mice having an acceptable signal-to-noise ratio were placed on a water regulation schedule, with provision of 99% hydrogel (HydroGel^®^, ClearH_2_O, USA) substituting for water intake. The daily amount of hydrogel was gradually decreased over a week from 2 g to the targeted amount. With this water restriction protocol, daily amounts were regulated to maintain body weight up to 85% of age- and sex-matched ad-lib group for the entire recording period. Based on weekly assessment of body condition scoring performed by veterinarian staff, an additional amount of hydrogel was added if necessary.

All mice had a one-day per week break from daily sessions. On the day before a break day, mice were provided with double size of their daily amount and on the break day, they resumed water restriction with their daily amount. In this manner, performance fluctuation often observed in ad-lib provision of water during a break could be minimized, and their overall health could be sustained.

### Apparatus

To maximize efficiency and throughput, we constructed 9 identical training apparatuses, each equipped with a fiber-photometry recording setup. Each apparatus housed a mounting plate and posts (Thorlabs) attached with small devices (head bar holder, reward delivery tube, photobeam sensor, light emitting diode (LED) panel, mouse body case holder, etc.), supported by 3D-printed support frames. Each apparatus was shielded with 0.5” thick soundproof sheets to minimize noise distractions during training. To minimize any mechanical sound generated by the solenoid controlling reward delivery, we hung the device outside of each training apparatus to avoid mice using the activating sound as an additional cue for reward delivery. All electronic devices in each apparatus were controlled by Arduino and Raspberry Pi systems, which generated timestamps of training events and TTL pulses that synchronized with time events for behavioral analysis. Each mouse was mounted on the apparatus by screws attached to their surgically attached head bar, and the small PVC tube otherwise kept in their home-cage was placed so as to encase their body during the training sessions. Each recording apparatus was threaded with a patch cord that delivered excitation and received emission signals connecting individually to integrated fluorescence Mini-cubes (Doric Lenses). Each fiber-coupled LED (405 and 470 nm, Thorlabs) was activated by an LED driver (Thorlabs) according to triggering pulses generated by a dual-channel multi-function waveform generator (Owon). The square pulses (1.5 msec) from each channel triggered 405 nm (isosbestic control) and 470 nm (green fluorescent protein (GFP) signal) LED drivers continuously at a rate of 30 Hz with the two pulses shifted 120° relative to each other. The emission signals were detected and amplified with a fluorescence detector (Doric Lenses). The excitation power of the LED driver was adjusted individually to achieve a peak emission signal intensity of 0.3 V for each excitation. This intensity was measured using a photodetector that converts fluorescent intensity to voltage. The detected signals as well as LED driving pulses and Arduino trial start pulse were acquired with a sampling rate of 10 kHz with T7-pro DAQ and LJStreamM software (LabJack).

### Training procedures

#### Sucrose solution habituation

On the first day of habituation, the reward spout delivering a 4% sucrose solution was placed close to the mouth of the head-fixed mouse and drops of solution were provided frequently until the mouse drank actively from the spout. The spout position was then moved gradually away from its mouth, requiring the mouse to protrude its tongue to lick the sucrose. Reward retrieval of a drop of solution (4 µL) then occurred by licking. Each tongue protrusion was detected as a lick event by a photobeam sensor installed at the side of the spout.

#### Lick-activity dependent and random reward habituation

After a mouse learned to retrieve the reward comfortably and actively, a lick activity-dependent (LAD) reward habituation session began. During this habituation session, when a lick event was detected, a drop of solution was given, followed by various intervals from 6 to 8 sec.

The 1-hr LAD habituation sessions continued daily until mice actively consumed more than 150 droplets per session. Most of the mice required 1-2 habituation sessions, but some mice required more sessions. Once mice exhibited active licking for reward consumption, they underwent random reward habituation sessions in which they received unexpected drops of reward with varying intervals of 8-48 sec, given for two or three sessions (**Fig. 6a**). Therefore, all mice underwent LAD reward habituation followed by random reward habituation before cue training sessions began. Random reward "probe" sessions were also inserted every 9 or 10 sessions during training on subsequent tasks to determine whether dopamine responses to random rewards would change over the longitudinal training sessions.

#### Single-cue and reward learning

Following random reward habituation, all mice began daily training sessions on visual cue and reward association. Water regulated mice were placed in the head-fixed apparatus, and a blue LED (intensity setting at 3 lux) was placed to present visual cues on the right and left side at eye level (**Fig. 1a**), and with the reward delivery/lick detection device placed near the mouth. Each trial started with an LED lit (cue) on one side, the cue was lit for 1.5 sec, the LED was turned off, and a reward was concurrently delivered. Each mouse had one of the two LEDs (right or left) designated as the cue predicting reward throughout training sessions. The locations of the reward cue were counterbalanced among subjects. One daily session typically consisted of ∼175 trials with randomly varying trial durations of 8-48 sec. The daily session continued until a mouse showed stable performance defined by greater than 0.75 in area under the receiver operating characteristic (auROC; see below) comparing lick counts during pre-cue and cue period for at least 2 consecutive sessions.

#### Cue discrimination and reversal discrimination training

After completion of single-cue and reward conditioning, daily cue discrimination training began by inserting, into the original schedule of rewarded trials, trials with the opposite LED presented without reward (non-rewarded trials). Therefore, for each training session, two trial types (rewarded and non-rewarded trials) were intermixed pseudo-randomly (with no more than a 4x sequential repetition of one trial type). Each trial started with either the left or right side LED lit (cue), and a reward was delivered 1.5 sec later at cue off only for the reward cue. Daily sessions, each typically consisting of around 350 trials with randomly varying trial duration of 8-12 sec, continued until the mouse exhibited a stable discrimination level defined by greater than 0.75 in auROC value between cue presentations for at least 2 consecutive sessions. Daily training sessions on reversal learning then began. The cue and reward contingencies were reversed. Mice learned that the previously rewarded cue was no longer rewarded, but that the previously non-rewarded cue now predicted an upcoming reward. Daily sessions of reversal learning continued until a mouse exhibited a stable discrimination defined by greater than 0.75 in auROC value between cue presentations for at least 2 consecutive sessions.

#### Probabilistic reward learning

After completion of reversal learning, probabilistic reward training proceeded. The rewarded cue given during the prior reversal learning was the only cue presented. For the daily probabilistic reward sessions, reward was provided partially according to the target probability and omitted reward trials were randomly selected for each session. Two sessions of reward probabilities of 0.75 and 0.5 were run, and for each reward probability, the session with the better recording quality was selected for analysis shown in **Fig. 3a and b**. A session with reward probability of 1.0 was always given before and after each probabilistic reward session. For some mice, two additional sessions having a block design of reward probabilities were performed, consisting of 4 blocks with different reward probabilities (0.75, 0.25, 0.5 and 1.0 reward probability with 44 trials per block) in a session.

#### Calculation of auROC

To evaluate a performance level for each session, lick numbers for the duration of interest for all trials were used to compute auROC as a learning index. For cue discrimination, lick numbers during 1.5 sec from cue onset were used and sorted according to trial types. For cue evoked licks, lick numbers were documented for the period from 1.5 sec before cue onset until cue onset (pre-cue period), and from cue onset until 1.5 sec after cue onset (cue period) for all cue presentations. To examine lick behavior for the reward cue, lick numbers from 1.5 sec before reward cue onset until reward cue onset (pre-reward cue period) and from 0 sec to 1.5 sec after reward cue onset (reward cue period) were separately tallied. The empirical auROC was calculated to represent a differential index of two datasets. The thresholds for constructing an auROC curve were taken for every middle point of all data sample differences of each given two data sets. Based on each threshold, true positive (TP; the number of incidents of one kind greater or equal to threshold; i.e., lick rate for rewarded trials or lick rate for cue period) and false negative (FN; the number of incidents of the same kind less than threshold) samples were computed to calculate TPR (true positive rate) as TP / (TP + FN) for each threshold. Similarly, false positive (FP; the number of incidents of the other kind greater or equal to threshold; i.e., lick rate for no rewarded trials or lick rate for pre-cue period) and true negative (TN; the number of incidents of the same kind less than threshold) were computed to calculate FPR (false positive rate) as FP / (FP + TN) for each threshold. The auROCs for cue discrimination (auROC_disc)_ or for cue evoked lick (auROC_evk_) were taken from the trapezoidal values of the ROC curves generated by FPR and TPR. The auROC_disc_ and auROC_evk_ were used as a learning index of session for cue discrimination (and reversal discrimination) and single-cue conditioning, respectively. The session auROC was mainly used to determine when to advance to the next learning schedule of each mouse. Each mouse was given up to 20 daily sessions, and if it failed to reach the learning criterion of at least 0.75 session auROC, the mouse was excluded from the following daily session schedules. Mice that reached the learning criterion were able to advance to the next phase of learning.

### Fiber-photometry recording

Each fiber-coupled LED (405 and 470 nm, Thorlabs) was activated by an LED driver (Thorlabs) according to triggering pulses generated by a dual-channel multi-function waveform generator (Owon). The square pulses (1.5 msec) from each channel triggered 405 nm (isosbestic control) and 470 nm (GFP signal) LED drivers continuously at a rate of 30 Hz, with the 470 nm pulse train lagging 120° behind to 405 nm pulse train. The emission signals were detected and amplified with a fluorescence detector (Doric Lenses). The emission signals from the fluorescence detector, TTL pulses to drive two LED drivers (Thorlab), and trial start TTL from the Arduino were acquired with a sampling rate of 10 kHz with a T7-pro DAQ and LJStreamM software (LabJack). To acquire emission signals from six mice in six separate apparatuses at one time, data acquisition of 6 analog inputs of the emission signal and 8 channels of TTL inputs (405 LED driver, 470 LED driver, and trial start TTLs of 6 Arduinos) was arranged through a CB37 terminal board (LabJack). Before each recording session, the excitation intensity was set to produce emission photodetector output around 0.3 V. Because the emission signal from 470 nm excitation fluctuated due to active GFP, the minimal amplitude was set to around 0.3 V. The digital inputs were then separated offline based on the connecting ports. The TTL for triggering two LED drivers was used to extract emission signals during the excitation period, and TTLs from each Arduino were matched to the corresponding analog channels. The last sample of each 1.5 msec pulse was taken as a reading value for the entire excitation pulse, and the emission samples were separated according to the excitation wavelength. The time of trial start TTL served as an event timestamp for the corresponding analog channels.

The data acquired from each session were prepared with several preprocessing steps. The raw data of a session for GFP (470 nm) and control (405 nm) signals were extracted and separated by excitation pulses and band-passed with a low pass filter (5 Hz). The max values for each GFP pulse were converted to (F_GFP_ − F_CNT_) / F_CNT_, where F_CNT_ denoted the max value obtained from the nearest control pulse (dF/F). Then, dF/F of low pass filtered data of each session data were z-score transformed.

### Statistical test with generalized linear mixed model

For statistical inferences, we used a generalized linear mixed model (GLMM) in R package (glmmTMB). Our main interest was to confirm the effects of recording region, learning type, and performance achieved on the dopamine release responses to cue and reward that were recorded. To quantify and build a model, we used AUC (trapezoidal method) of the dopamine trace (z transformed dF/F data) for the rewarded trials. The trial trace was prepared by calibrating with its own baseline (subtracting mean of 1.5-sec pre-cue value from each data point), and trial AUC of early cue (0.5-sec period from cue onset), late cue (0.5-sec period to cue offset), and reward (1-sec period from the first lick after reward) and were used as response variables. For testing the effects of sex on dopamine response, we tested the rewarded trial data from cue conditioning and cue discrimination (with regressors learning kind x sex x region on early and late cue and reward). For estimating the effect of performance on the dopamine responses, we examined the rewarded trial data from the last session of cue conditioning, cue discrimination, and reversal discrimination (learning kind x performance level x region).

### Discrimination performance level

The performance level was determined by the auROC value at the last session and its group median for discrimination between rewarded and non-rewarded trials based on lick counts during the cue presentation period. A mouse having a higher auROC value than group median on the last session was assigned as a "better" (higher) performer, otherwise the mouse was assigned as a "worse" (lower) performer. The interaction model was chosen over an additive model based on the ANOVA likelihood ratio test of two models. The post-estimation process of GLMM results then was performed with the emmeans package in R to check the effect of learning kinds over other levels (learning, region, sex, or performance).

The initiation of spontaneous licks was detected if a lick occurred during the ITI period (period starting 6.5 sec after cue onset and ending with the next cue onset) and was preceded by a period without licks of at least 0.5 sec. Lick-aligned fluorescence trace data were prepared based on each detected lick (0.5-sec pre-lick and 1-sec post-lick data), and each lick trace was calibrated using the pre-lick data as baseline. The AUC of lick traces was obtained for 0.5-sec period after each lick. These lick AUCs were used as response variables on learning kinds, learning and region in GLMM model and post-estimation.

### Histological localization of recording sites

We defined four reference points in each coronal section: the most medial point of striatum (x1, y1), the most lateral point of striatum (x2, y2), the most dorsal point of striatum (x3, y3), and the most dorsal point of anterior commissure (x4, y4). Designating the tip of probe as (x5, y5), the standardized medial-lateral coordinate of the probe tip was calculated as (x5 – x1) / (x2 – x1), and the standardized depth coordinate of the probe tip was calculated as (y5 – y3) / (y4 – y3). Recording sites were classified as "medial" if the distance from the midline to the tip of the probe was less than 0.6 of the distance from the midline to the lateral edge of the striatum.

### Anatomical maps of first principal component

Principal components analysis calculations were performed by the Matlab ‘pca’ function, using each mouse’s waveform of dF/F, averaged over all rewarded trials in that mouse’s final ("acquisition") session of discrimination training, as input. Spatial smoothing of the PC1 amplitude values across mice was done in each cardinal anatomical plane as follows. First a coordinate grid of 50 bins was constructed to span the relative position values in each direction, resulting in bins whose widths depended on the 3D axes (0.012 wide in the M-L direction, 0.016 in the D-V direction, 0.012 in the A-P direction). For each mouse, the value of the amplitude ("score") for PC1 was assigned to the bin containing the recording site’s coordinates and was copied throughout a square of 17×17 neighboring bins extending eight bins to each side of the bin containing the recording site, truncated if necessary to stay within the bounds of the 50×50 coordinate grid. All other bins were assigned the value NaN. Each mouse produced a single set of 50×50 bins. Two computations were than performed across the 63 sets of bins corresponding to the 63 mice: the total number of non-NaN values was counted for each bin position, and the NaN-tolerant average value was computed for each bin position (i.e., the mean obtained strictly from the mice that had non-NaN-valued bins at a given position, or NaN if there were no non-NaN values). The average values of any bins where the total number of non-NaN values in the bin was less than 3 were then reset to NaN, resulting in a 50×50 matrix where every bin contained either NaN or the average of values from at least 3 mice. The 50×50 matrix was then plotted as a pseudo-color image, with the color scale chosen so that its limit in the negative direction, representing NaN, was well below the most negative value in any bin.

To assess statistical significance of the spatially smoothed average values, we performed bootstraps on the set of mice included in the entire calculation. Two-hundred bootstraps were performed by randomly selecting 63 mice at a time, with replacement (i.e., a given mouse could be repeated) from the actual set of 63 mice recorded. The median value across bootstraps was used to create the final pseudo-color plots, and the median of those median values was used as a reference value for statistical significance. If the 2.5th percentile of values across bootstraps was greater than the reference value, that bin was marked as significantly high. If the 97.5th percentile of values was less than the reference value, the bin was marked significantly low.

## Acknowledgments

We thank Dr. Steven Worthington for consulting and helping on statistical approach, Henry F. Hall for constructing recording apparatus, Dr. Sebastien Delcasso for designing the apparatus system, Dr. Yasuo Kubota for help with manuscript preparation, Dr. Ayano Matsushima for reading a pre-final version of the manuscript, and Jonny Loftus for help with figure preparation. This work was funded by the National Institutes of Health (R01 MH060379), the William N. & Bernice E. Bumpus Foundation (RRDA Pilot: 2013.1), the Saks Kavanaugh Foundation, the CHDI Foundation (A-5552), and Dr. Lisa Yang.

## Author contributions

A.M.G. and M.J.K. designed and initiated the research; M.J.K. and D.H. performed the surgeries; M.J.K, C.S., P.S. and K.T. performed the recordings; D.H., M.J.K, C.S. performed perfusion; A.M and T.Y performed the histology and imaging; D.H., C.S., P.S., and K.T provided animal handling and care; M.J.K. and D.J.G. analyzed data with detailed A.M.G input; A.M.G., D.J.G., and M.J.K wrote the manuscript.

## References

1 Schultz, W., Dayan, P. & Montague, P. R. A neural substrate of prediction and reward. Science 275, 1593–1599 (1997).

2 Sutton, R. S. & Barto, A. G. Reinforcement Learning: An Introduction. First Edition edn, (MIT Press, 1998).

3 Romo, R. & Schultz, W. Dopamine neurons of the monkey midbrain: contingencies of responses to active touch during self-initiated arm movements. J Neurophysiol 63, 592–606 (1990).

4 Joshua, M., Adler, A., Mitelman, R., Vaadia, E. & Bergman, H. Midbrain dopaminergic neurons and striatal cholinergic interneurons encode the difference between reward and aversive events at different epochs of probabilistic classical conditioning trials. J Neurosci 28, 11673–11684 (2008).

5 Cohen, J. Y., Haesler, S., Vong, L., Lowell, B. B. & Uchida, N. Neuron-type-specific signals for reward and punishment in the ventral tegmental area. Nature 482, 85–88 (2012).

6 Puryear, C. B., Kim, M. J. & Mizumori, S. J. Conjunctive encoding of movement and reward by ventral tegmental area neurons in the freely navigating rodent. Behav Neurosci 124, 234–247 (2010).

7 Eshel, N. et al. Arithmetic and local circuitry underlying dopamine prediction errors. Nature 525, 243–246 (2015).

8 Robinson, S., Sandstrom, S. M., Denenberg, V. H. & Palmiter, R. D. Distinguishing whether dopamine regulates liking, wanting, and/or learning about rewards. Behav Neurosci 119, 5–15 (2005).

9 Starkweather, C. K., Babayan, B. M., Uchida, N. & Gershman, S. J. Dopamine reward prediction errors reflect hidden-state inference across time. Nat Neurosci 20, 581–589 (2017).

10 Berke, J. D. What does dopamine mean? Nat Neurosci 21, 787–793 (2018).

11 Lerner, T. N. et al. Intact-brain analyses reveal distinct information carried by SNc dopamine subcircuits. Cell 162, 635–647 (2015).

12 Howe, M. W. & Dombeck, D. A. Rapid signalling in distinct dopaminergic axons during locomotion and reward. Nature 535, 505–510 (2016).

13 Parker, N. F. et al. Reward and choice encoding in terminals of midbrain dopamine neurons depends on striatal target. Nat Neurosci 19, 845–854 (2016).

14 Patriarchi, T. et al. Ultrafast neuronal imaging of dopamine dynamics with designed genetically encoded sensors. Science 360, eaat4422 (2018).

15 Sun, F. et al. A genetically encoded fluorescent sensor enables rapid and specific detection of dopamine in flies, fish, and mice. Cell 174, 481–496 e419 (2018).

16 Liu, C. et al. An action potential initiation mechanism in distal axons for the control of dopamine release. Science 375, 1378–1385 (2022).

17 Threlfell, S. et al. Striatal dopamine release is triggered by synchronized activity in cholinergic interneurons. Neuron 75, 58–64 (2012).

18 Brimblecombe, K. R. & Cragg, S. J. The striosome and matrix compartments of the striatum: a path through the labyrinth from neurochemistry toward function. ACS Chem Neurosci 8, 235–242 (2017).

19 Hamid, A. A., Frank, M. J. & Moore, C. I. Wave-like dopamine dynamics as a mechanism for spatiotemporal credit assignment. Cell 184, 2733–2749 e2716 (2021).

20 Krok, A. C., Mistry, P., Li, Y. & Tritsch, N. X. Intrinsic reward-like dopamine and acetylcholine dynamics in striatum. bioRxiv https://doi.org/10.1101/2022.09.09.507300 (2022).

21 Cox, J. & Witten, I. B. Striatal circuits for reward learning and decision-making. Nat Rev Neurosci 20, 482–494 (2019).

22 Saunders, B. T., Richard, J. M., Margolis, E. B. & Janak, P. H. Dopamine neurons create Pavlovian conditioned stimuli with circuit-defined motivational properties. Nat Neurosci 21, 1072–1083 (2018).

23 Tsutsui-Kimura, I. et al. Distinct temporal difference error signals in dopamine axons in three regions of the striatum in a decision-making task. Elife 9 (2020).

24 Hikosaka, O., Kim, H. F., Yasuda, M. & Yamamoto, S. Basal ganglia circuits for reward value-guided behavior. Annu Rev Neurosci 37, 289–306 (2014).

25 Choi, K. et al. Distributed processing for action control by prelimbic circuits targeting anterior-posterior dorsal striatal subregions. bioRxiv https://doi.org/10.1101/2021.12.01.469698 (2021).

26 Choi, K., Holly, E. N., Davatolhagh, M. F., Beier, K. T. & Fuccillo, M. V. Integrated anatomical and physiological mapping of striatal afferent projections. Eur J Neurosci 49, 623–636 (2019).

27. Matsumoto, M. & Hikosaka, O. Two types of dopamine neuron distinctly convey positive and negative motivational signals. Nature 459, 837–841 (2009).

28 Bromberg-Martin, E. S., Matsumoto, M. & Hikosaka, O. Dopamine in motivational control: rewarding, aversive, and alerting. Neuron 68, 815–834 (2010).

29 Brischoux, F., Chakraborty, S., Brierley, D. I. & Ungless, M. A. Phasic excitation of dopamine neurons in ventral VTA by noxious stimuli. Proc Natl Acad Sci U S A 106, 4894–4899 (2009).

30 Markowitz, J. E. et al. Spontaneous behaviour is structured by reinforcement without explicit reward. Nature 614, 108–117 (2023).

31 Dai, B. et al. Responses and functions of dopamine in nucleus accumbens core during social behaviors. Cell Rep 40, 111246 (2022).

32 Howe, M. W., Tierney, P. L., Sandberg, S. G., Phillips, P. E. & Graybiel, A. M. Prolonged dopamine signalling in striatum signals proximity and value of distant rewards. Nature 500, 575–579 (2013).

33 Prager, E. M. et al. Dopamine oppositely modulates state transitions in striosome and matrix direct pathway striatal spiny neurons. Neuron 108, 1091–1102 e1095 (2020).

34 Nadel, J. A. et al. Optogenetic stimulation of striatal patches modifies habit formation and inhibits dopamine release. Sci Rep 11, 19847 (2021).

35 Sgobio, C. et al. Aldehyde dehydrogenase 1-positive nigrostriatal dopaminergic fibers exhibit distinct projection pattern and dopamine release dynamics at mouse dorsal striatum. Sci Rep 7, 5283 (2017).

36 Graybiel, A. M. & Matsushima, A. The ups and downs of the striatum: Dopamine biases upstate balance of striosomes and matrix. Neuron 108, 1013–1015 (2020).

37 Jeong, H. et al. Mesolimbic dopamine release conveys causal associations. Science 378, eabq6740 (2022).

38 Coddington, L. T., Lindo, S. E. & Dudman, J. T. Mesolimbic dopamine adapts the rate of learning from action. Nature 614, 294–302 (2023).

39 Cone, I., Clopath, C. & Shouval, H. Z. Learning to express reward prediction error-like dopaminergic activity requires plastic representations of time. bioRxiv https://doi.org/10.1101/2022.04.06.487298 (2023).

40 Amo, R. et al. A gradual temporal shift of dopamine responses mirrors the progression of temporal difference error in machine learning. Nat Neurosci 25, 1082–1092 (2022).

41 Akiti, K. et al. Striatal dopamine explains novelty-induced behavioral dynamics and individual variability in threat prediction. Neuron 110, 3789–3804 e3789 (2022).

42 Takahashi, Y. K. et al. Dopaminergic prediction errors in the ventral tegmental area reflect a multithreaded predictive model. Nat Neurosci 26, 830–839 (2023).

43 Hamid, A. A. et al. Mesolimbic dopamine signals the value of work. Nat Neurosci 19, 117–126 (2016).

44 Mohebi, A. et al. Dissociable dopamine dynamics for learning and motivation. Nature 570, 65–70 (2019).

45 Lee, R. S., Engelhard, B., Witten, I. B. & Daw, N. D. A vector reward prediction error model explains dopaminergic heterogeneity. bioRxiv https://doi.org/10.1101/2022.02.28.482379 (2022).

46 Berridge, K. C. & Robinson, T. E. What is the role of dopamine in reward: hedonic impact, reward learning, or incentive salience? Brain Res Brain Res Rev 28, 309–369 (1998).

47 Lee, R. S., Mattar, M. G., Parker, N. F., Witten, I. B. & Daw, N. D. Reward prediction error does not explain movement selectivity in DMS-projecting dopamine neurons. Elife 8 (2019).

48 Glowinski, J., Cheramy, A., Romo, R. & Barbeito, L. Presynaptic regulation of dopaminergic transmission in the striatum. Cell Mol Neurobiol 8, 7–17 (1988).

49 Cragg, S. J. & Greenfield, S. A. Differential autoreceptor control of somatodendritic and axon terminal dopamine release in substantia nigra, ventral tegmental area, and striatum. J Neurosci 17, 5738–5746 (1997).

50 Nelson, A. B. et al. Striatal cholinergic interneurons drive GABA release from dopamine terminals. Neuron 82, 63–70 (2014).

51 Beatty, J. A., Song, S. C. & Wilson, C. J. Cell-type-specific resonances shape the responses of striatal neurons to synaptic input. J Neurophysiol 113, 688–700 (2015).

52 Thorn, C. A. & Graybiel, A. M. Differential entrainment and learning-related dynamics of spike and local field potential activity in the sensorimotor and associative striatum. J Neurosci 34, 2845–2859 (2014).

53 Wilson, C. J. Predicting the response of striatal spiny neurons to sinusoidal input. J Neurophysiol 118, 855–873 (2017).

54 Graybiel, A. M. & Matsushima, A. Striosomes and Matrisomes: Scaffolds for Dynamic Coupling of Volition and Action. Annu Rev Neurosci 46, 359–380 (2023).

55 Vu, M. T. et al. in International Basal Ganglia Society Meeting.

