## Supplementary Figure 1 for "Dopamine Release Plateau and Outcome Signals in Dorsal Striatum Contrast with Classic Reinforcement Learning Formulations"

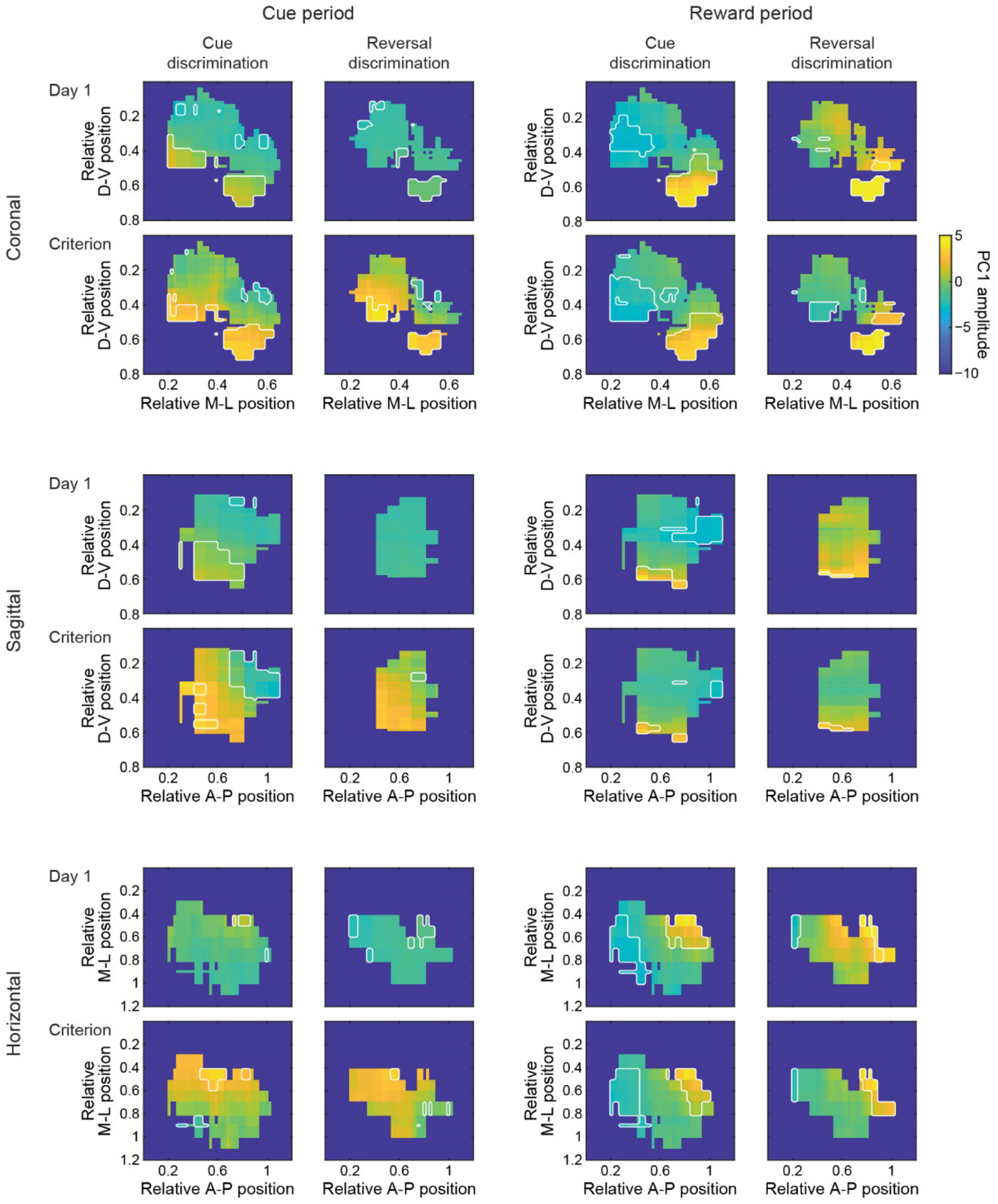


**Supplementary Fig. 1**. Anatomical distribution of PC1 amplitude for the first (Day 1) and last (Criterion) sessions in cue discrimination and cue reversal training, in all three orthographic projection planes. For this analysis, all the waveforms of four sets of sessions (cue discrimination day 1, cue discrimination criterion, cue reversal day 1, cue reversal criterion) were analyzed together in one pass of PCA, to make the PC amplitudes directly comparable. The first 3 PC waveforms were similar to those seen when analyzing only the Cue Discrimination − Criterion waveforms (**Fig. 5**), as were the qualitative patterns of the PC1 anatomical maps (amplitudes differed slightly in value). Reduction in reward PC1 amplitude with learning was only seen in the dorsal-most sites, and only in reversal training.
